# Global Scientific Trends in Organoids from 2004 to 2023: A Data-Driven Bibliometric and Visualized Analysis

**DOI:** 10.1101/2024.08.28.610094

**Authors:** Min Zhao, Liangju Kuang, Haoxin Guo, Xindan Cao, Junshi Dai, Yupeng Wang, Zhongqing Wang, Cheng Peng

## Abstract

To conduct a bibliometric analysis of organoids to describe international research trends and visualize current research directions. This cross-sectional bibliometric analysis examined the development of organoid research from 2004 to 2023. The current study used VOS-viewer to assess and analyze 13,174 documents. Literature data were collected on a specific date (Feb 19, 2024) and downloaded in plain text from Web of Science Core Collection. In this paper, 13,174 organoid papers were retrieved from Web of Science Core Collection. There were only 114 organoid studies in 2004, and from 2015 onward, the number of annual publications on this topic began to proliferate, reaching 10,023 from 2019 to 2023, accounting for as much as 76.1% of the total number of published papers. The United States proudly leads the way in both the volume of articles published and the number of citations garnered, standing tall as the undisputed frontrunner. Among the illustrious institutions, Harvard University and the University of Washington are among the most prolific. Hans Clevers has worked with 121 prolific authors and has the most publications. With the use of organoids in cancer modeling, drug screening, and regenerative medicine, organoid technology has attracted much attention in medicine, and the significant increase in the number of published papers and citations signifies the expanding influence and global collaboration in the field of organoid research. This study contributes to our understanding of current trends and potential future advances in the field of organoid research by identifying five distinct clusters in the field.

## 1. Introduction

Organoids[1] are self-organizing three-dimensional tissues, typically induced from pluripotent and adult stem cells, capable of recreating in vitro the structure and function of in vivo organoid tissues, mimicking the critical functions and biological complexity of organs, and capable of being cultured in a stable, long-term succession. Organoid models are therefore considered to be superior models in the detection of human diseases compared to two-dimensional cell cultures and animal models, and organoid technology holds great promise in the field of cancer biology[2]. Organoid models are more amenable to manipulating ecotope components and signaling pathways and genome editing than traditional two-dimensional cultures and animal models. In addition, organoids can be combined with gene editing technology[3, 4], microfluidics[5, 6], nanotechnology[7, 8], and artificial intelligence[9], which not only expands the scope of application but also promotes the development of organoid technology. Collaborative interdisciplinary research[10–16] in organoid technology is driving innovation in life sciences and medicine, contributing to the advancement of human health and medical technology.

Organoid technology is gradually developing and maturing, and the relevant literature is increasing annually. However, few reviews have comprehensively analyzed the literature related to organoids, and the purpose of this paper is to compensate for this deficiency and provide a panoramic view of organoids for subsequent researchers. This study aimed to utilize bibliometrics and visualization techniques to analyze organoid research publications from the last two decades, thereby revealing trends, identifying key influencers, exploring collaboration networks, and pinpointing research hotspots and emerging themes.

## 2. Method

### 2.1 Data collection

The original data for this study were retrieved from the Web of Science (WOS) database, which is a comprehensive literature database comprising multiple indices such as the SCIE, SSCI, A&HCI, and CPCI. Altogether, it houses over one billion searchable citations spanning more than 250 disciplines, making it a globally recognized and authoritative source. We utilized the Science Citation Index Expanded (SCIE) database within the Web of Science Core Collection for data collection using the following search query: Topic= (“Organoid$” or “Microphysiological System$” or “Organ Chip$” or “Organ* on Chip$” or “Organ*-on-a-Chip$” or “Organotypic Cell Culture$” or “Organotypic Culture$” or “Organotypic Model$”). The time frame for publications was set between January 1, 2004, and December 31, 2023, and non-English language publications were excluded. The document types were limited to articles and reviews. All relevant bibliometric information was downloaded on February 19, 2024, to minimize the impact of database updates. A summary of the data sources and selection criteria is shown in Figure 1.

**Figure 1.**
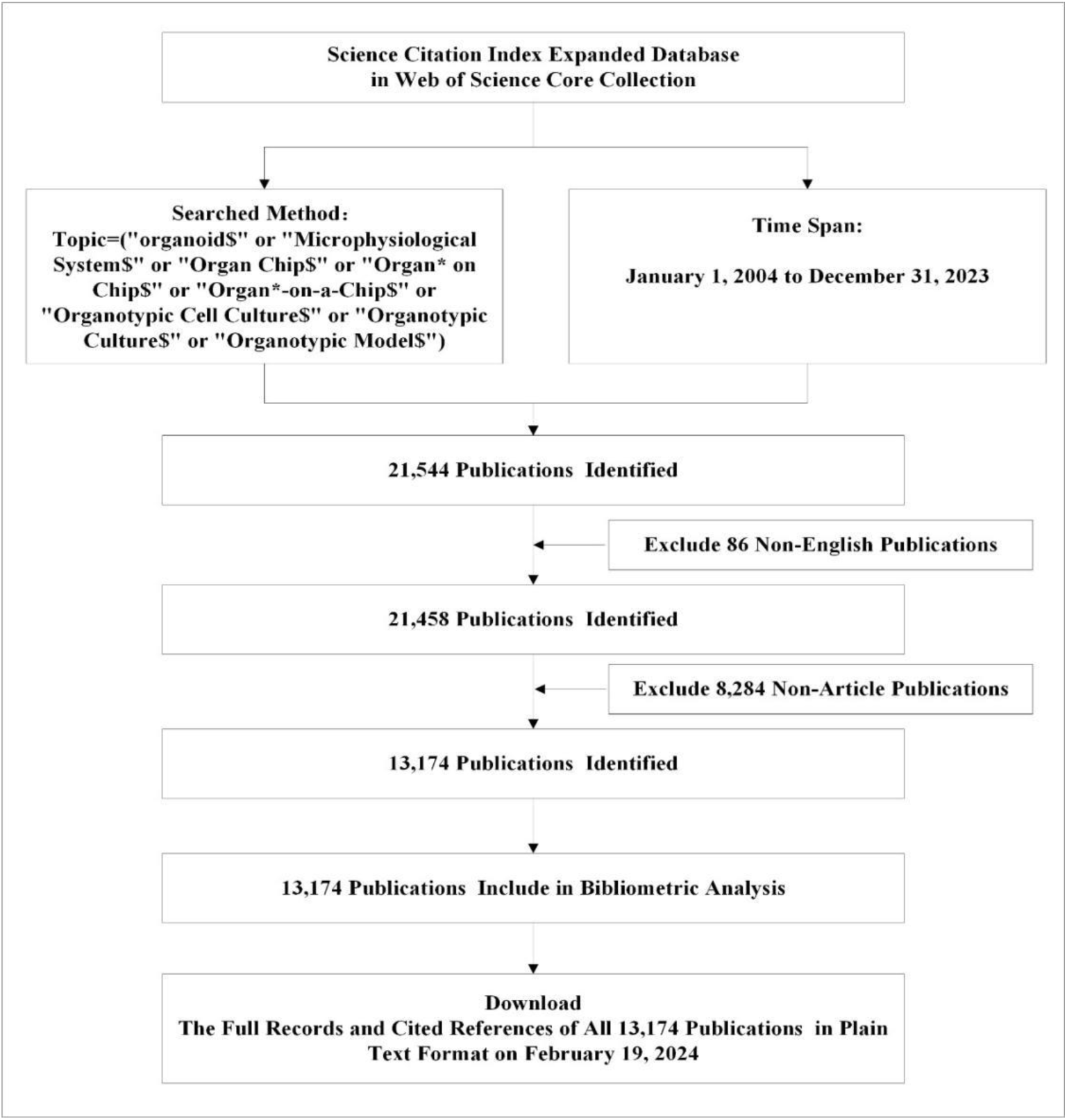
Detailed flowchart steps of the search strategy for screening literature.

### 2.2 Analysis tools and visualization maps

VOSviewer[17] is a widely used bibliometric analysis software that facilitates various types of qualitative studies. In this study, the analysis was carried out using VOSviewer 1.6.20. Co-authorship analyses were performed for institutions and authors, co-citation analyses were carried out for references, and co-occurrence analyses were carried out for keywords. In addition, we performed descriptive analyses for the year of publication, journal, country, institution, and author.

## 3. Results

### 3.1 Publishing trends

The trend of publication quantity from 2004 to 2023 is shown in Figure 2, with a total of 13174 organoid publications retrieved. There were only 114 organoid studies in 2004, which exceeded 1000 for the first time in 2019 and peaked at 2,367 in 2023. The number of publications in 2023 is 20.76 times that in 2004. From 2004 to 2014, the yearly number of publications for this research was reasonably stable with slight fluctuations, remaining between 114 and 235 and showing a steady growth trend. Beginning in 2015, the number of annual publications started to increase, and the rate of growth accelerated each year. The number of publications from 2019 to 2023 reached 10023, accounting for as much as 76% of the total number of publications. The number of publications on organoid studies between 2004 and 2023 shows a trend from steady growth to explosive growth.

**Figure 2.**
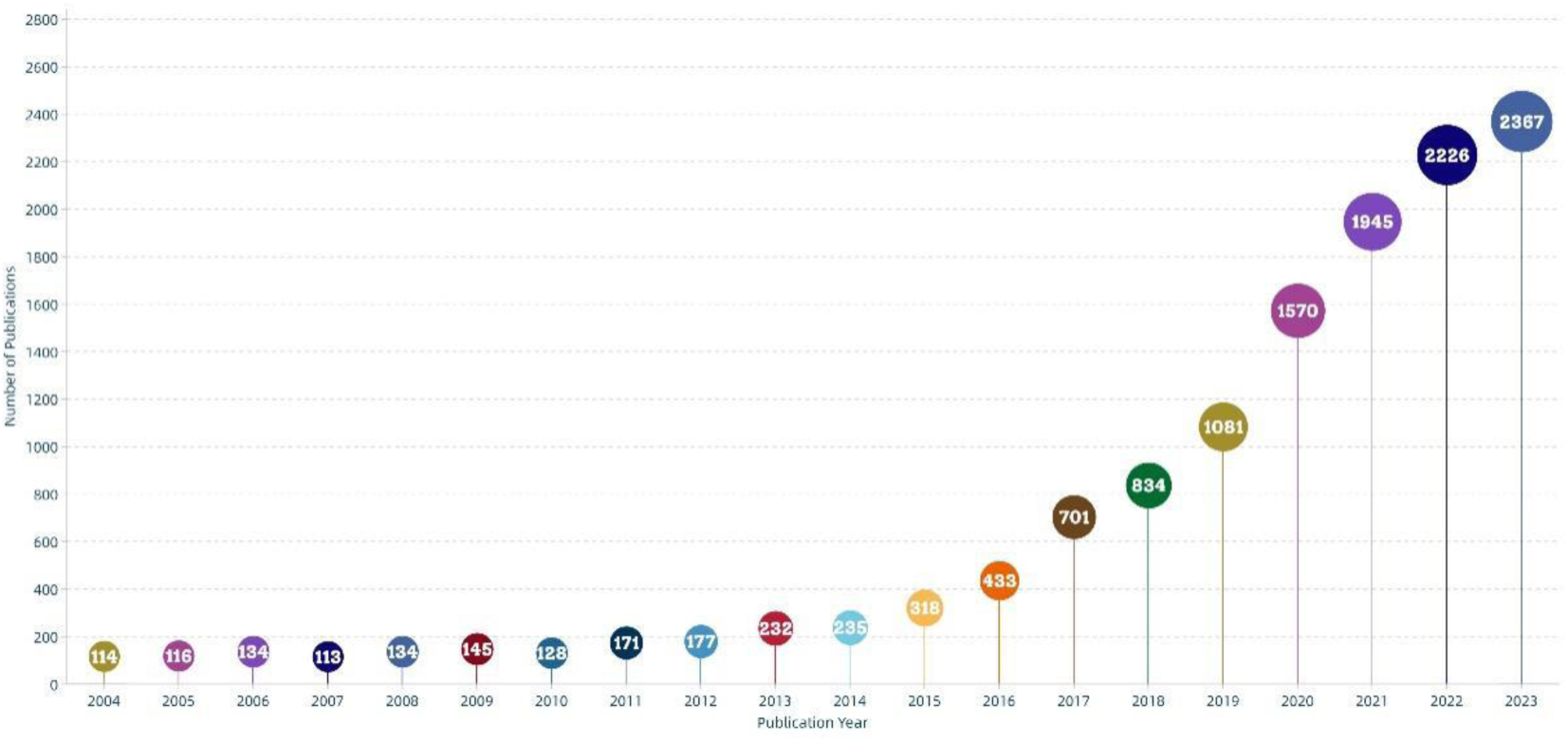
Trends in publications from 2004 to 2023.

### 3.2 Publishing trends

In total, 1,587 journals have published groundbreaking research on organoids. By establishing a threshold of 30 publications, we narrowed down to 86 journals that have significantly contributed to the field, accounting for 6,588 publications, representing a substantial 50% of all organoid-focused works. To further explore these influential journals, we employed VOSviewer to conduct a citation analysis. The resulting overlay visualization map in Figure 3 portrays each journal’s impact, with sphere dimensions representing the number of publications and a color gradient from blue to red indicating the average citation count, ranked from lowest to highest. Our findings indicate that Scientific Reports led with 400 publications, followed by Nature Communications with 298, and the International Journal of Molecular Sciences with 226. Notably, fifteen journals achieved an average citation count exceeding 50, with Cell topping the list as the most highly cited journal, closely followed by Nature and Nature Medicine.

**Figure 3.**
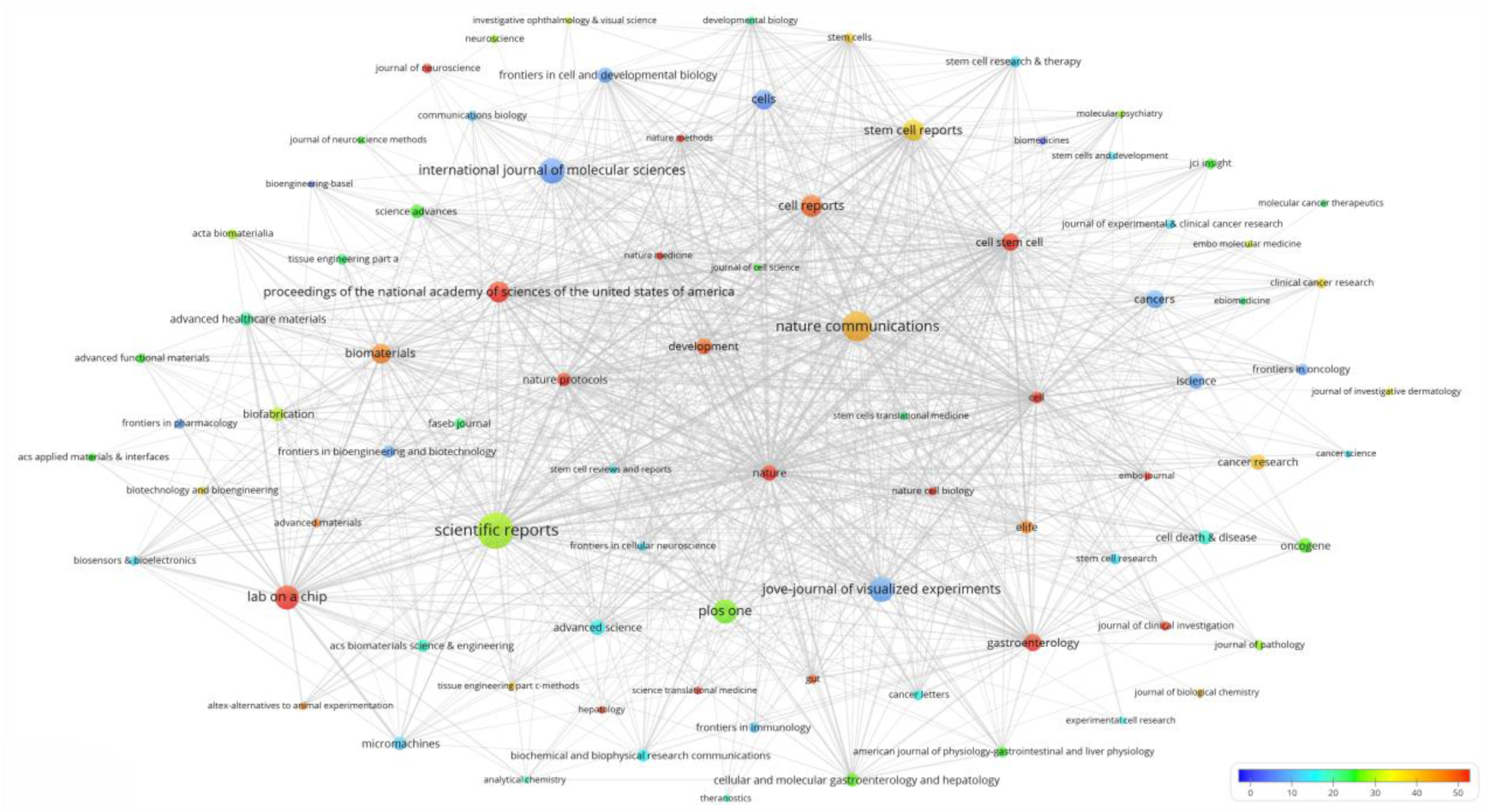
Network map of high-yield journal coauthor ship.

### 3.3 Country/Region

Globally, 138 countries/regions are involved in organoid research, and their global distribution is shown in Figure 4. There were 28 countries/regions with more than 100 publications. There were 6 countries with more than 1,000 publications: including two from North America, two from Europe, and one each from Asia and Oceania. The top ten countries, according to the number of publications, are displayed in Table 1, with the United States (6,413 counts, 192,616 citations) ranking first concerning both publications and citations. The United Kingdom (1,679 counts, 46,253 citations), China (1,386 counts, 21,629 citations), and Germany (1,372 counts, 39,992 citations) were the following top countries. The Netherlands had the greatest mean citations among the top ten countries, while China had the highest average publication year.

**Figure 4.**
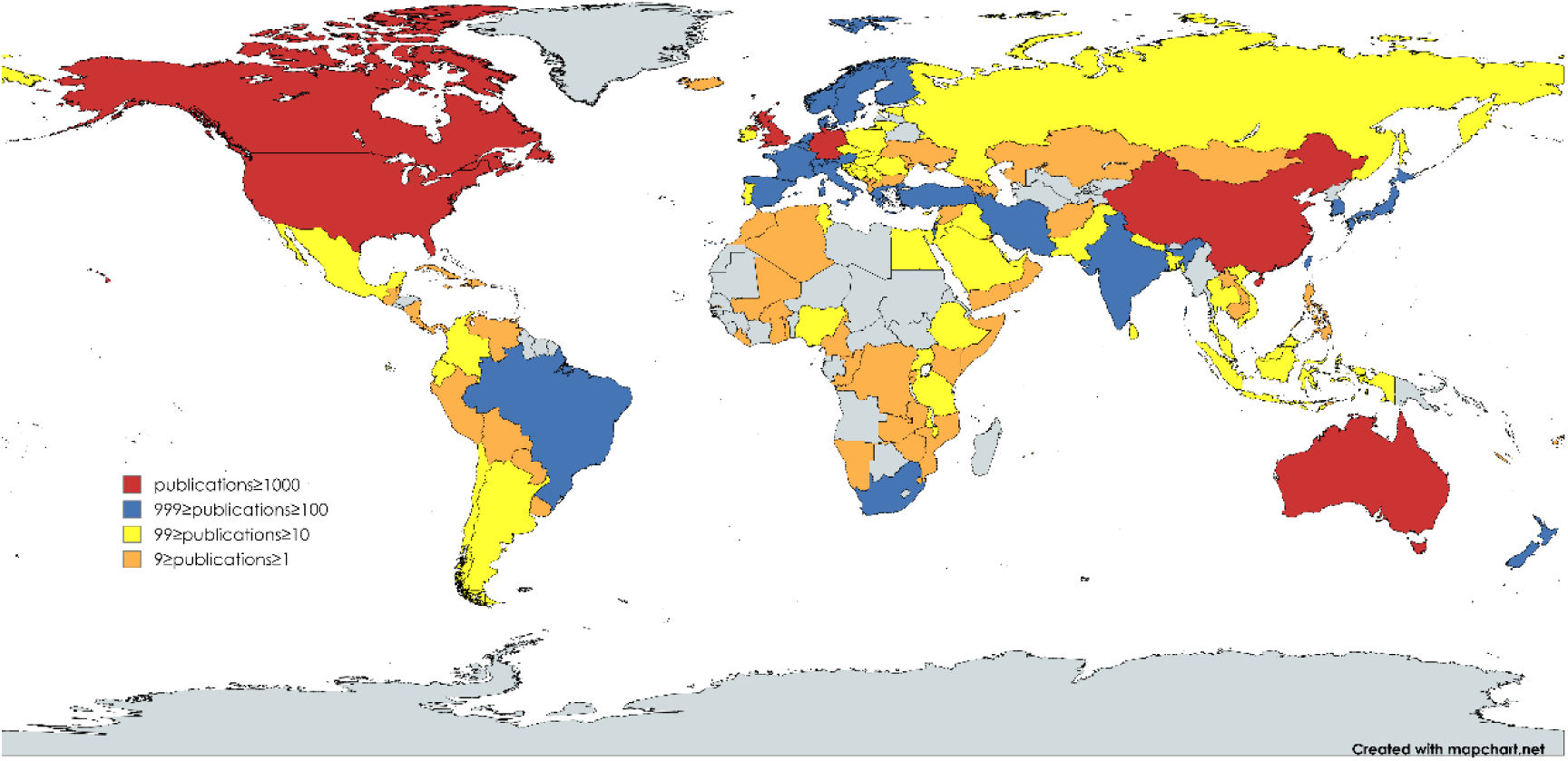
Global spatial distribution of the number of articles published.

**Table 1.** Top 10 countries by number of publications.

| Rank | Country | Counts | Citations | Avg. Citations | Avg. Pub. Year |
| --- | --- | --- | --- | --- | --- |
| 1 | United States of America | 6413 | 192616 | 30 | 2014.47 |
| 2 | United Kingdom | 1679 | 46253 | 28 | 2015.55 |
| 3 | China | 1386 | 21629 | 16 | 2017.01 |
| 4 | Germany | 1372 | 39992 | 29 | 2014.70 |
| 5 | Australia | 1314 | 34065 | 26 | 2014.87 |
| 6 | Canada | 1002 | 34190 | 34 | 2014.68 |
| 7 | France | 606 | 20211 | 33 | 2015.12 |
| 8 | Italy | 600 | 17622 | 29 | 2014.53 |
| 9 | Netherlands | 579 | 20119 | 35 | 2015.20 |
| 10 | Switzerland | 480 | 13453 | 28 | 2015.52 |

### 3.4 High Institutions

There are 9331 institutions involved in organoid research publications, with the ten most prolific institutions detailed in Table 2. Harvard University (n=544) boasted a publication account and emerged as the leader in this field, trailed closely by the University of Washington (n=284), the University of Chinese Academy of Science (n=254), and Johns Hopkins University(n=238). Among the top 10 institutions, Harvard University (n=44,924) has the most citations, followed by the University of Washington(n=13,026). Among the top 10 institutions, University Medical Center Utrecht (n=171) had the most citations. Among the top 10 institutions, the University of Chinese Academy of Science had the newest average publication year. We set the threshold for institutions to publish 30 times and identified 260 highly productive institutions out of 9,331 institutions. Using VOSviewer, we conducted a co-authorship analysis on these 260 high-producing institutions, revealing a robust institutional co-authorship network comprising 7 distinct clusters, as depicted in Figure 5

**Figure 5.**
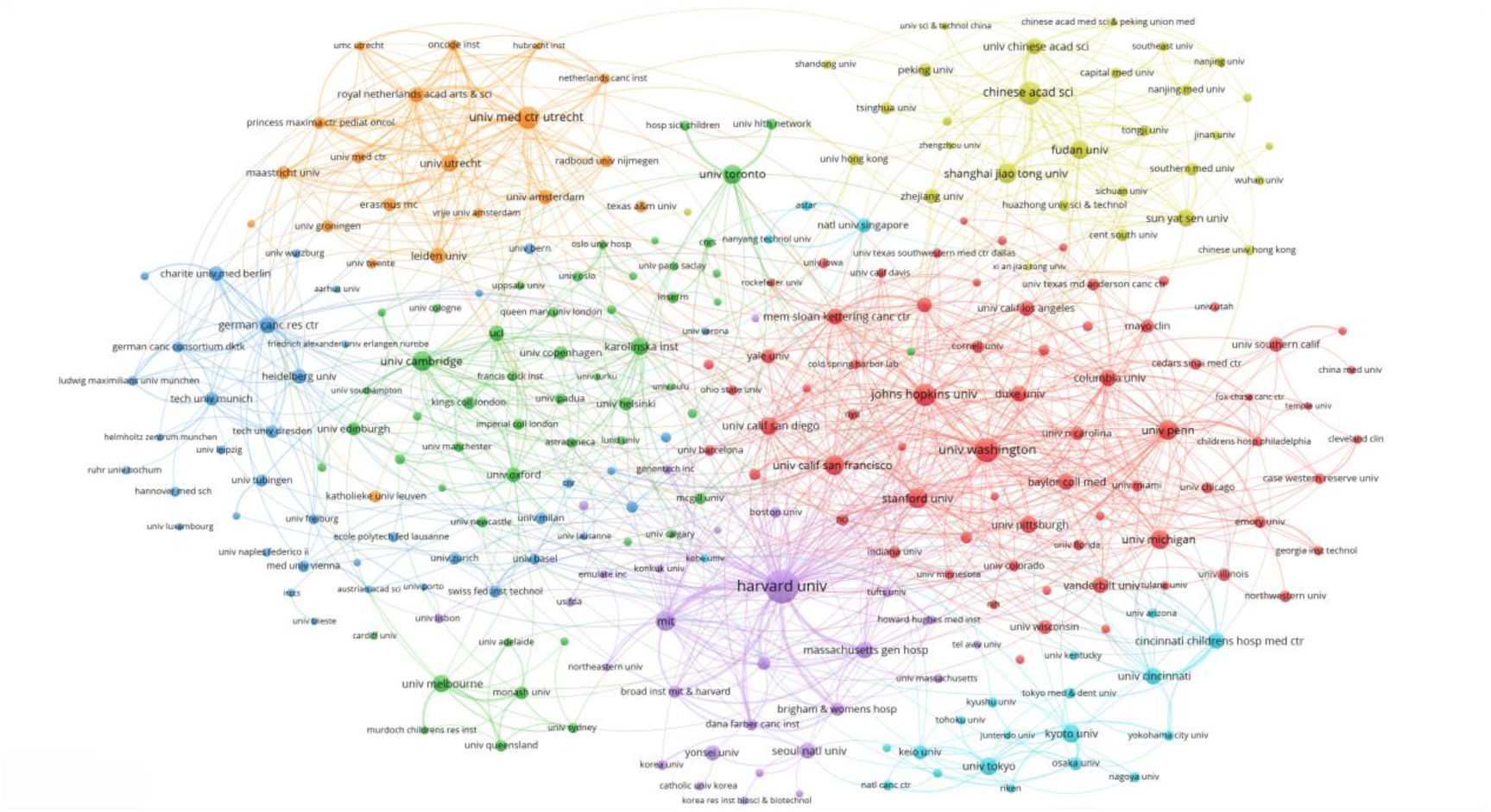
Highly productive institutional co-authorship network map.

**Table 2.** Top 10 institutions with the most publications.

| Rank | Institution | Counts | Citations | Avg. Citations | Avg. Pub. Year |
| --- | --- | --- | --- | --- | --- |
| 1 | Harvard University | 544 | 44924 | 83 | 2019.09 |
| 2 | University of Washington | 284 | 13016 | 46 | 2019.16 |
| 3 | University of Chinese Academy of Science | 254 | 6393 | 25 | 2020.98 |
| 4 | Johns Hopkins University | 238 | 15525 | 65 | 2019.10 |
| 5 | University Medical Center Utrecht | 237 | 40427 | 171 | 2018.71 |
| 6 | Massachusetts Institute of Technology | 193 | 19935 | 103 | 2019.13 |
| 7 | University of California, San Francisco | 191 | 12183 | 64 | 2019.06 |
| 8 | Stanford University | 189 | 11798 | 62 | 2019.28 |
| 9 | University of Pennsylvania | 189 | 12627 | 67 | 2018.28 |
| 10 | University of Cambridge | 184 | 14988 | 81 | 2019.35 |

The largest cluster is represented by a vibrant red, encompassing 69 institutions in total. At the forefront is Harvard University, which has collaborative research ties with 219 highly productive institutions. Close behind is the University of Washington, which has partnered with 162 such institutions. Stanford University also boasts a strong presence in this network, with ties to 162 high-producing institutions, while the University of Pennsylvania has connections with 59 such entities.

### 3.5 Authors

A total of 84,733 authors contributed to organoid publications, and the top 12 authors by number of publications are listed in Table 3. At the forefront stands Clevers, Hans (n=220) having the most publications, trailed closely by Spence, Jason R. and Van Der Laan, Luc J. W. Among the top 12 authors, Clevers, Hans (n= 47,752) stands out as the most cited author, followed by Spence, Jason R.(n=6425). Among the top 12 authors, Sato, Toshiro (n=379) was the most cited on average, Van Der Laan, Luc J. W., and Verstegen, Monique M. A. were the newest in terms of average publication year. We set the threshold of authors posting 10 times and identified 429 exceptionally productive authors from 84,733 authors. We conducted a co-authorship analysis of the 429 high-producing authors using VOSviewer, which is presented in Figure 6.

**Figure 6.**
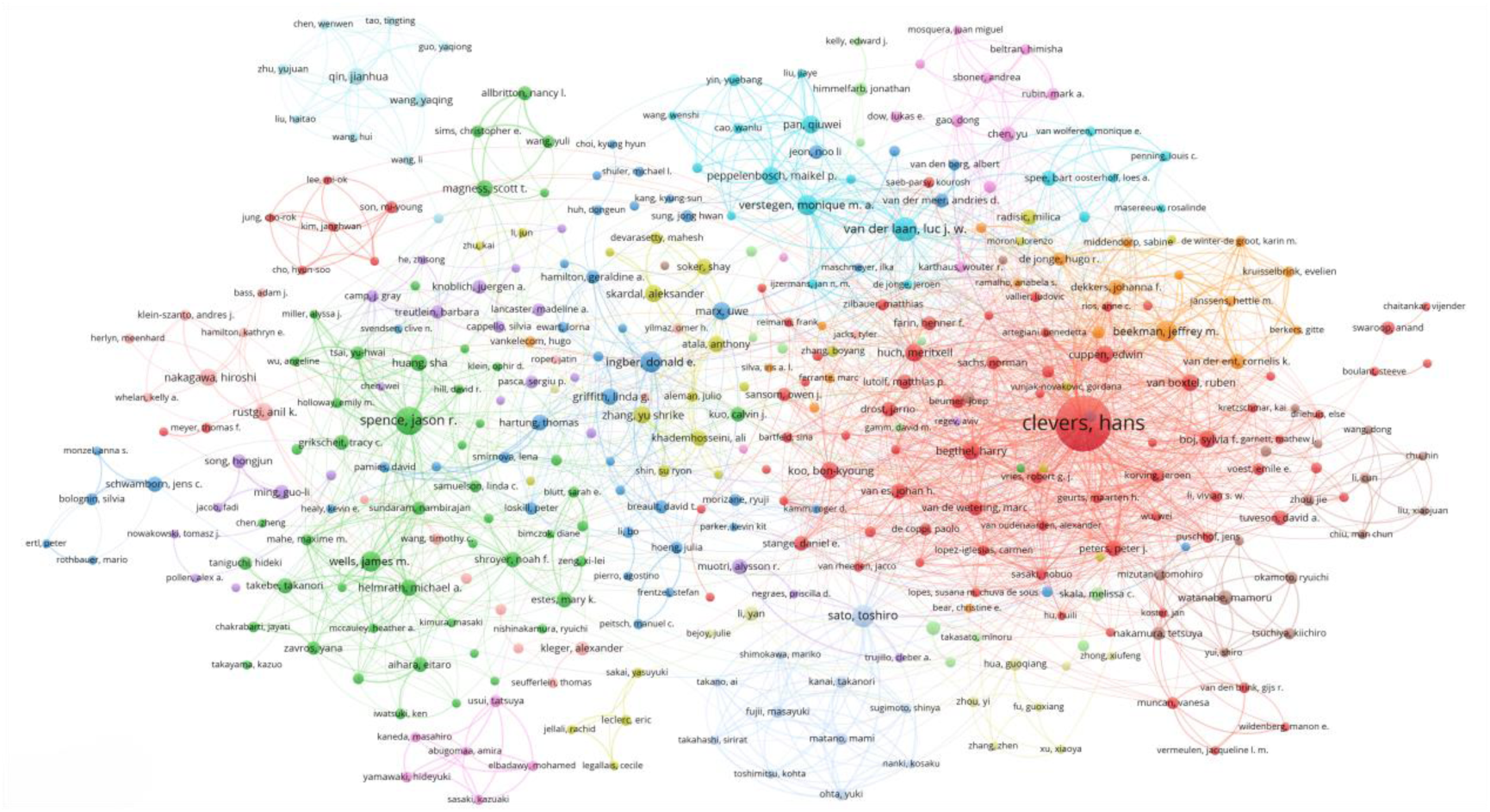
Prolific author co-authorship network map.

**Table 3.**
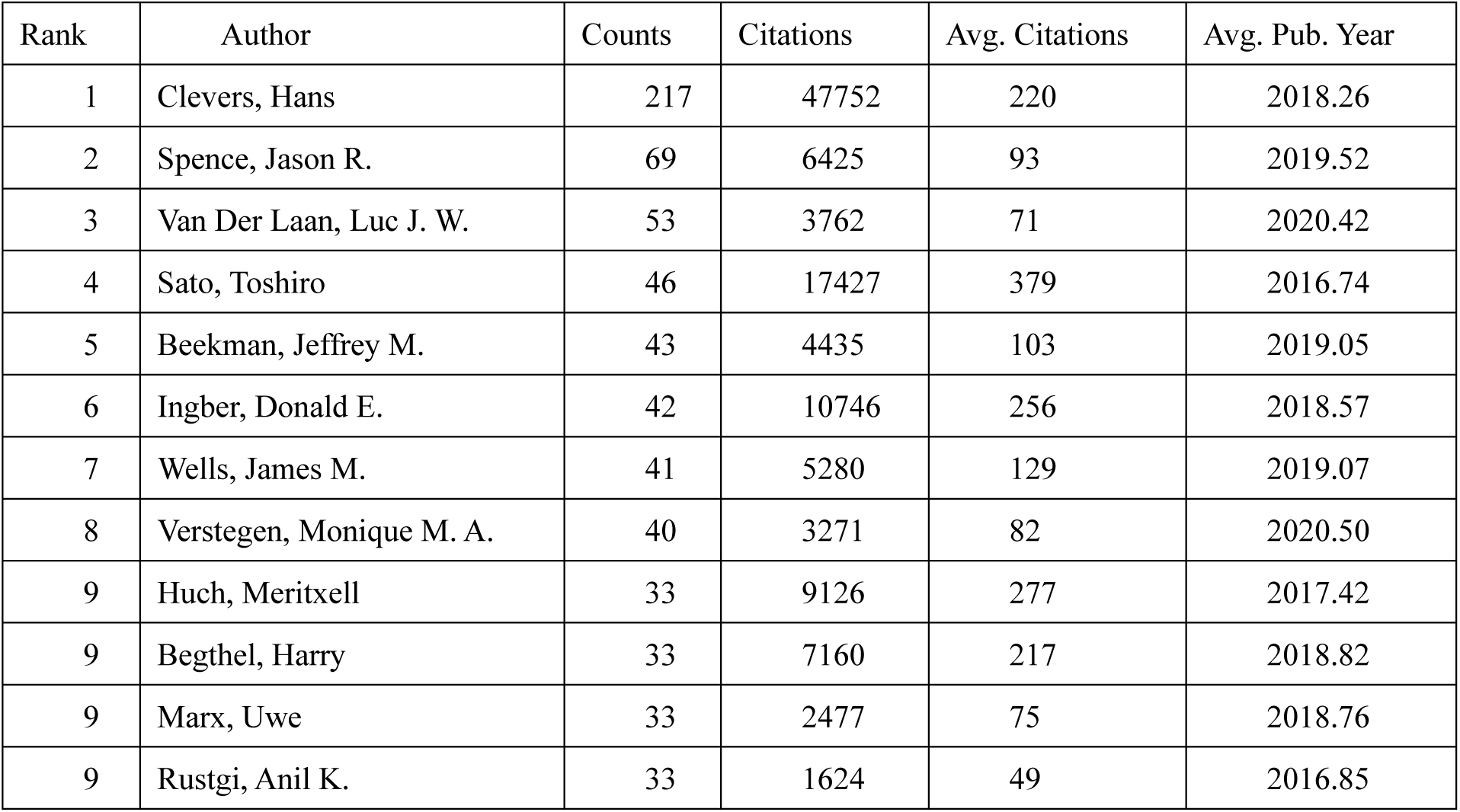
Top 12 authors of publications.

Of the 429 prolific authors, 369 formed the largest co-authorship network, encompassing 14 clusters. Hans Clevers has forged partnerships with 121 prolific authors. Edwin Cuppen has collaborated with 57 prolific authors, such as van Boxtel, and Ruben, who have collaborated with 56 prolific authors. Van Der Laan, Luc J. W., and Verstegen, Monique M. A., who collaborated 40 times, stand as the most closely aligned collaborators in the esteemed field.

### 3.6 References

Scientific research is carried out based on previous research, and analyzing references can help us understand the knowledge base of a particular area of study. There were 384,803 references cited, a total of 689,461 times from 13,174 organoid publications. On average, 52 references were cited per publication. We set the citation threshold of references to 100 and selected 162 highly cited references from the 384,803 references. We used VOSviewer to analyze and map the co-citations of these 162 highly cited references, as shown in Figure 7. As shown in the figure, Sato’s two articles[12, 18] from 2009 and 2011 are the most highly cited, in which he used adult stem cells from the mouse intestine to grow the first miniature intestinal organoids, which opened up the era of organoid research. The following highly cited paper is Lancaster’s 2013 paper[19], which recreates human development and disease through brain organoids. This highly cited literature has a high degree of influence and recognition in the field and provides a solid foundation for subsequent research.

**Figure 7.**
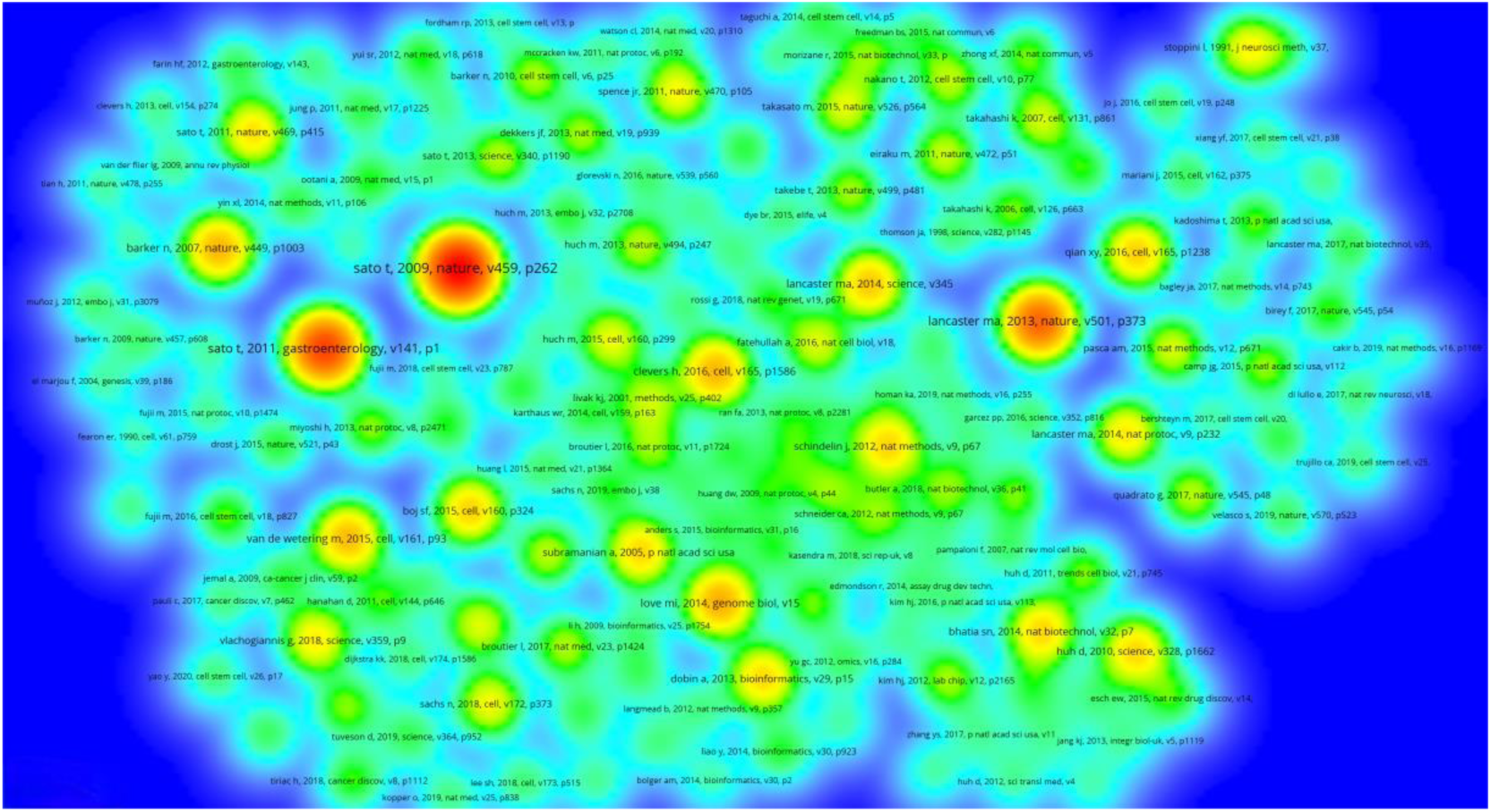
Co-citation density map of highly cited references.

**Figure 8.**
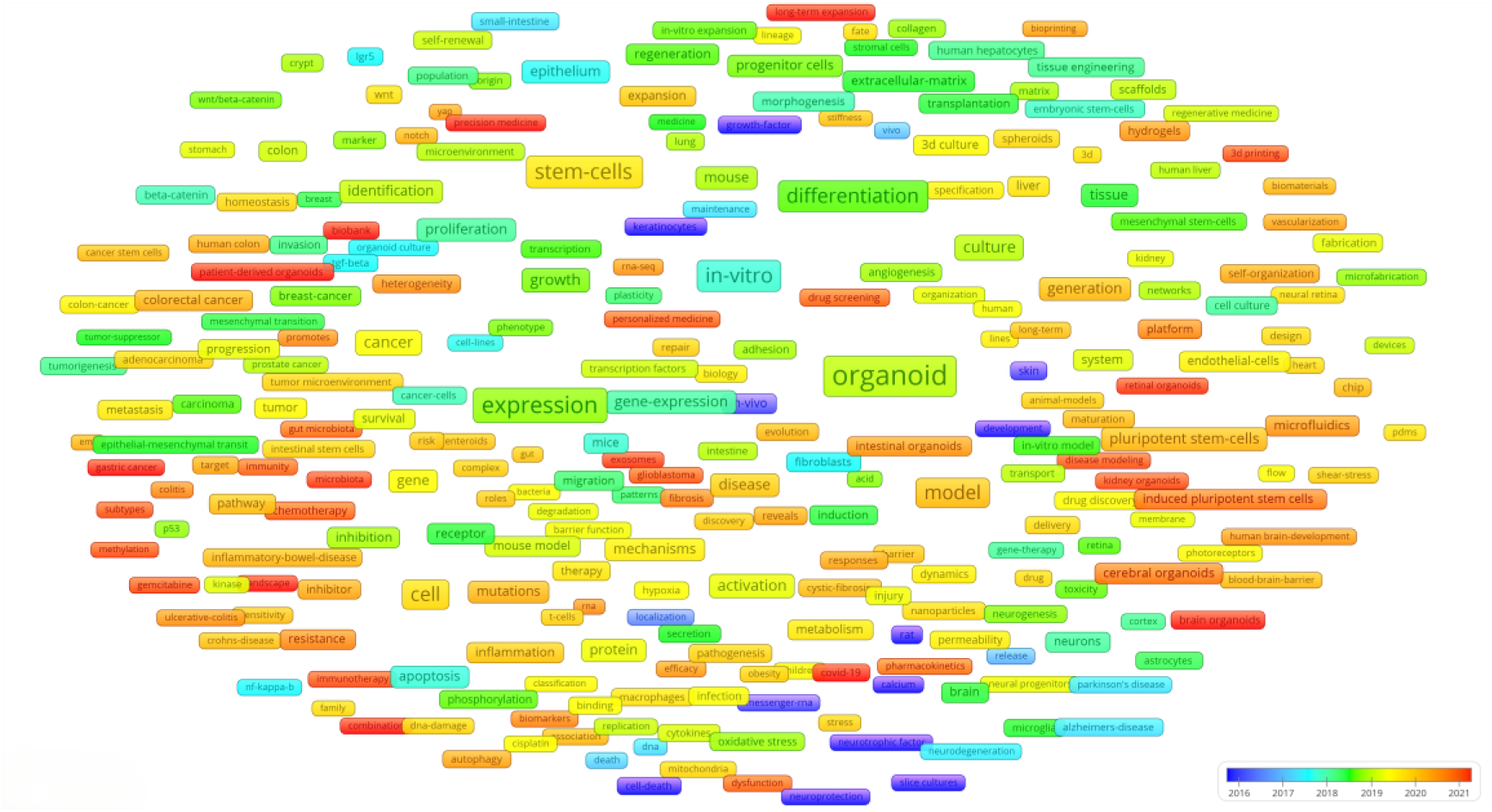
Co-occurrence network diagram of highly frequent keywords.

### 3.7 Keywords

There were 33,915 keywords in 13,174 articles, and we set the threshold for high-frequency keywords to 50, identifying 312 high-frequency keywords. Co-occurrence analysis was performed on the 312 keywords, and a co-occurrence network map of the high-frequency keywords was constructed, as shown in Figure 9. In addition to the search keyword class organ, expression (n=2,286), stem cells (n=1,838), in vitro (n=1,629), differentiation (n=1,554), and model (n=1,406) were the five keywords with the highest frequency of occurrence.

**Figure 9.**
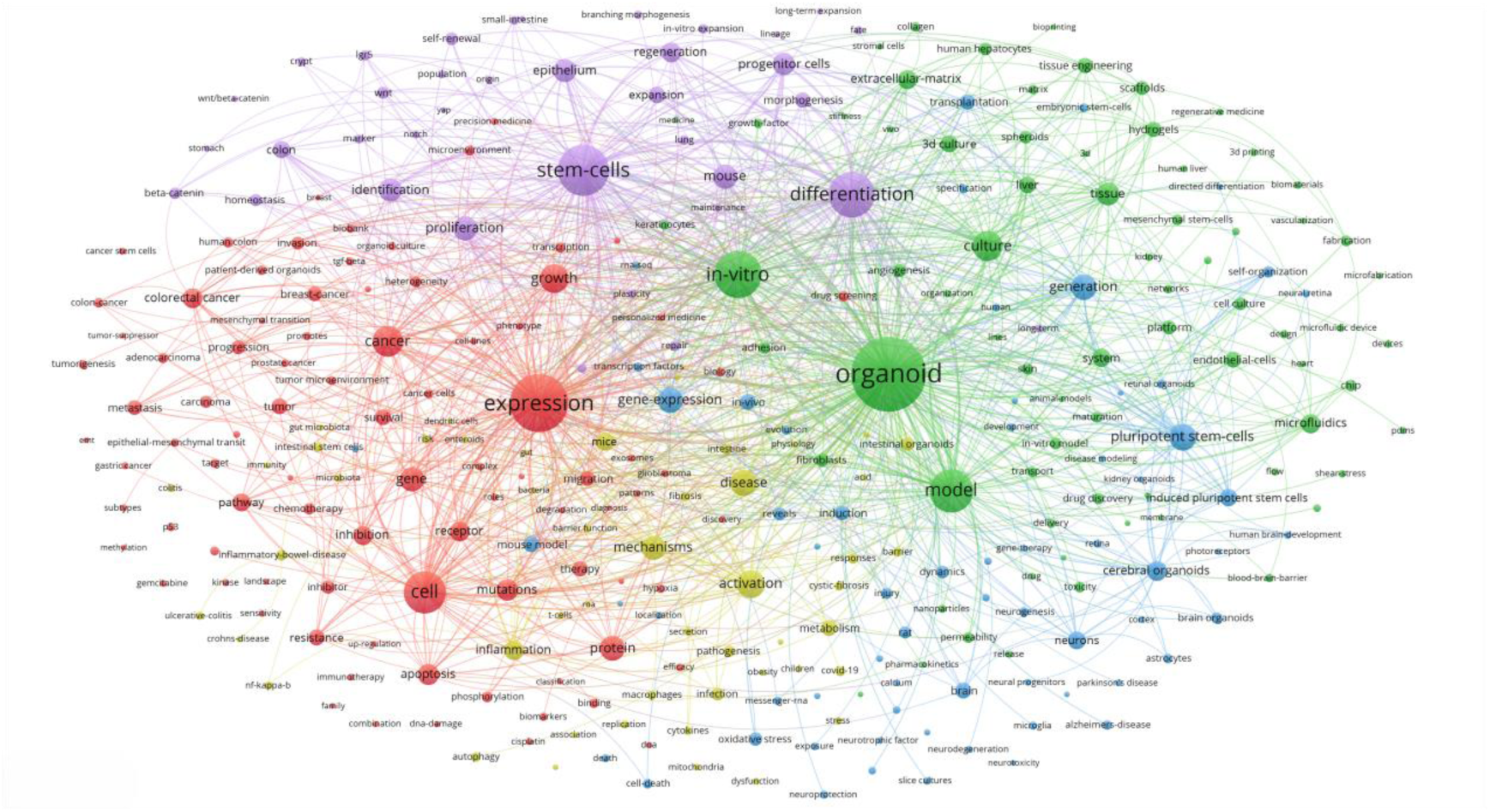
Co-occurrence overlay graph for highly frequent keywords.

We also constructed overlay graphs for high-frequency keywords, as shown in Figure 10. Co-occurrence analysis of high-frequency keywords revealed five clusters represented by different colors. Cluster 1, which focuses on organoid disease models, is red. The red cluster consists of 88 keywords and is the largest cluster. Cluster 2 is green and focuses on the role of organoids in drug screening. Cluster 3 is blue and describes the technology used for organoid culture. Cluster 4 is yellow, which focuses on the use of organoid models to explain disease-causing mechanisms at the cellular and molecular levels, and cluster 5 is purple, which describes the application of organoids in regenerative medicine.

**Figure 10.**
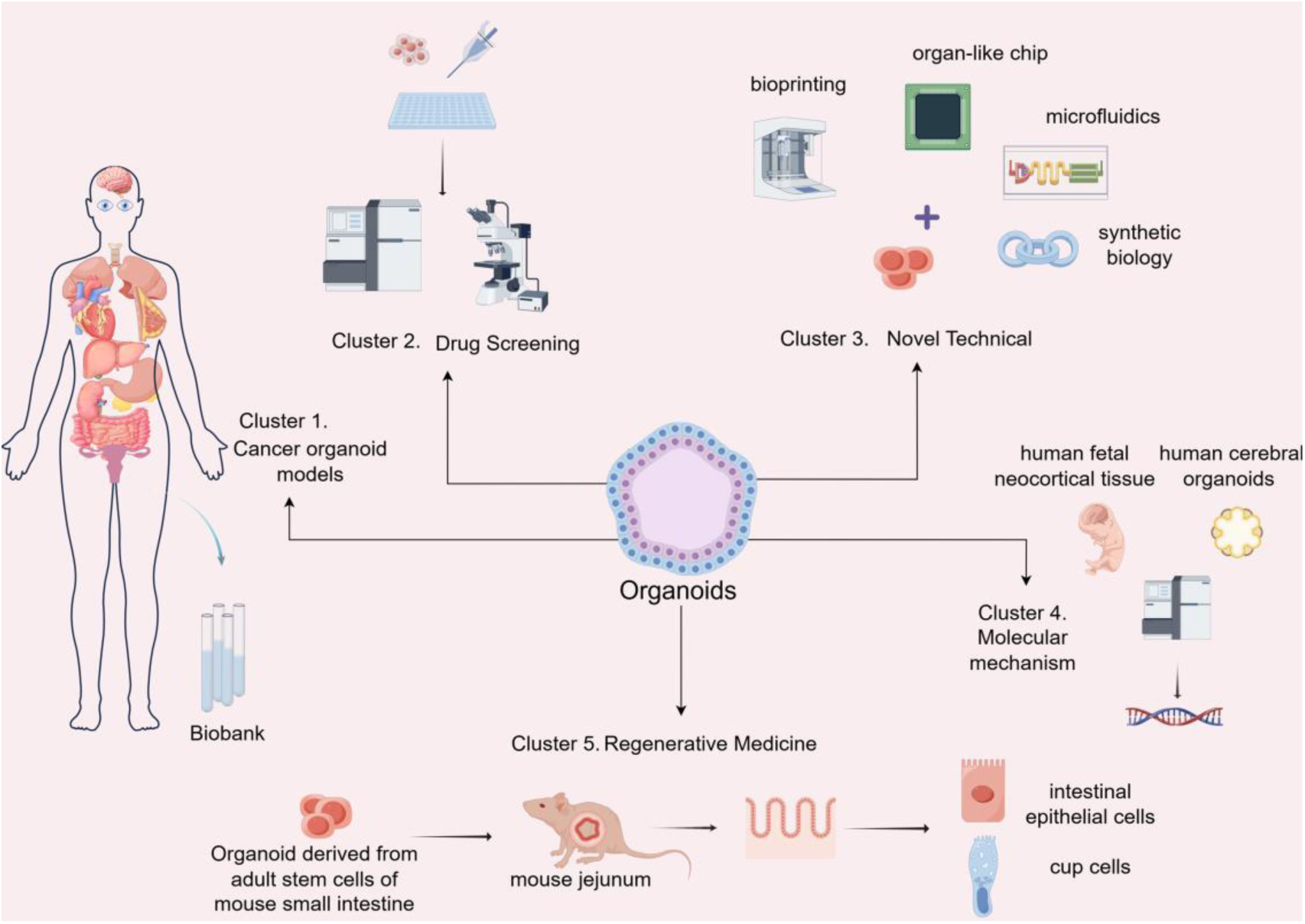
Major research hotpots around organoids.

The color of each node in the graph represents the average publication year of that node, with increasingly more recent publication rangeing from blue to red.

## 4. Discussion

Organoid research is a rapidly developing field with a strong international presence led by the United States in terms of research output and impact. The U.K., China, and Germany also play significant roles, while the Netherlands and China excel in citation impact and research recency. This bibliometric analysis revealed that the U.S. has the greatest number of publications, with Harvard University collaborating with 219 institutions, making it a global hub for collaboration alongside other top universities. As a revolutionary scientific research tool, organoid technology mimics the functions and properties of real organs by using three-dimensional structures cultured from human stem cells. This provides a powerful resource for various fields, including medical research, new drug development, regenerative medicine, and personalized treatment. In the future, the impact of organoid technology is expected to increase even further, playing a crucial role in advancing human health and medical technology.

### 4.1 Publishing trends

Our analysis of organoid research from 2004 to 2023 revealed a significant increase in the number of publications related to organoids, with most of the publications occurring in the last five years of the study period (Fig 3). The rapid growth in the field indicates a growing interest in and importance of organoid research in the scientific community. The high number of publications in journals, as well as the wide involvement of countries and regions worldwide, highlights the global impact and collaborative nature of organoid research. The United States dominates the field in terms of the number of publications and citations, which could be attributed to the country’s robust research infrastructure, funding, and expertise in the field. However, the Netherlands stands out as a leader in terms of the average number of citations per article, indicating a high level of influence and quality of research in the country (Fig 5).

### 4.2 International cooperation

The collaboration between top-performing institutions and authors has facilitated the advanced and widespread use of organoid applications in medical research. Leading universities such as Harvard, the University of Washington, Stanford, and the University of Pennsylvania act as centers for international cooperation, integrating diverse expertise and technology through interdisciplinarity for the progress of organoid technology (Fig 6). Cooperation between high-productivity institutions promotes academic exchange between institutions, sharing of technologies and resources, and translational cooperation between research results all contribute to accelerating the speed of scientific development while providing a broader perspective and more possibilities for organoid research on a global scale.

### 4.3 Research hotpots and emerging themes

#### 4.3.1 Cluster 1. Cancer organoid models

Tumor heterogeneity[20] due to environmental and genetic factors is a major challenge in current research on tumor biology. Compared with animal models and cellular models, organoids can reproduce the structure and function of in vivo tissues in vitro. Patient-derived tumor organoids can mimic the characteristics of in vivo tumors and the heterogeneity of tumor cells[21], which can provide new ideas for personalized cancer therapies and preclinical models for human disease research. In 2009, Hans Clever cultivated the first mouse-derived miniature intestinal organoids[12], followed by Sato et al., who used organoid technology to establish tumor organoids[18]. With the development of organoid culture technology, researchers have successfully cultured many cancer organoids, including rectal[22], gastric[23, 24], pancreatic[25, 26], breast[27, 28], and glioblastoma[29, 30]. By using tumor cells from specific patients to culture organoids, researchers can study individual differences in cancer in greater depth and provide a basis for precision medicine and personalized treatment strategies. Many biobanks of living organoids[27, 31–33] have been established, and large biobanks of healthy and tumor-like organs can be used to simulate the diversity of patient populations observed in large-scale studies. The Netherlands, the United States, and China are the countries with the most significant numbers of biobanks.

The construction of cancer-like organs consists of two main steps: human tumor tissue containing pluripotent, adult, or cancer stem cells is dissociated into small fragments or single cells using mechanical or chemical methods and then cultured in hydrogels with extracellular matrix components under appropriate culture conditions. Culture conditions vary for different tumor organoids, such as the composition of the medium, the addition of growth factors and hormones, and the oxygen partial pressure and carbon dioxide concentration. In addition, specific cultural techniques and physical conditions also vary.

The lack of standardization and technical differences in culturing tumor-like organs affects the determination of tumor heterogeneity and limits their preclinical application. The key issue in cancer organoid models is the lack of a cancer microenvironment, including stromal and immune cells. In recent years, with the advancement of research technologies, both source specimen collection and culture environments have gradually become standardized, and in addition to culture platforms, single-cell sequencing technology, microfluidics, and CRISPR-Cas9 gene editing can also be used to reduce the heterogeneity of tumor organoids. The three major difficulties in organoid research, namely “vascularization,” “immunization,” and “systemization,” have been solved by different teams[34–36] in recent years.

#### 4.3.2 Cluster 2. Drug screening

Currently, the cost of new drug development is high, the clinical translation rate is low[37], and conventional cell lines or animal models have certain limitations[38]. Most promising clinical drugs have failed to be effectively utilized in human therapy and cell lines differ greatly from those used in the human environment, tumors are highly heterogeneous, and it is difficult for animal models to accurately mimic human development and disease due to the limitations of biological differences. Patient-derived tumor-like organs retain the morphological and genetic properties and typical biomarker expression of the original tumor tissues and tumor heterogeneity, based on which they can be stably passed on, making them ideal disease-specific models for high-throughput screening of drugs. Tissue-derived tumor-like organoid models, which are usually constructed using the matrix gel method, are currently more frequently used for screening drugs. The constructed model will be validated morphologically, histologically, and molecular genetically, and multiple fluorescence immunohistochemistry and single-cell sequencing methods can be used to determine whether the organoid is consistent with the original tumor tissue.

Drugs that can be screened in organoids include chemotherapeutic drugs, small molecule targeted drugs, and antibody drugs. Hans Clevers’ team established a model of nonalcoholic fatty liver disease[39], screened for effective drug candidates for steatosis, and revealed the molecular mechanisms by which drugs inhibit lipogenesis. Herpers et al.[40] conducted a high-throughput screen of dual-targeted bispecific antibodies against organoid biobanks derived from colorectal cancer patients and paired healthy colonic mucosal samples and identified a bispecific antibody, called MCLA-158, that blocks the developmental and survival pathways of cancer cells by degrading EGFR proteins in cancer stem cells containing the LGR5 marker. Pellegrini[41] established choroid plexus-like organs that secrete cerebrospinal fluid and mimic the human blood-brain barrier to qualitatively and quantitatively predict the permeability of new drugs and selectively regulate the entry of small molecules such as dopamine and levodopa, which can be used as a screening platform for evaluating predicted human blood-cerebrospinal fluid drug permeability.

The use of various organoids[42–45] to test potential drug candidates allows for more accurate prediction of the effects of drugs in the human body, thus increasing the efficiency of drug development. As preclinical models, organoids are also promising for drug toxicity testing and efficacy evaluation[46], which can provide a reference for late-stage clinical drug use.

#### 4.3.3 Cluster 3. Novel Technical

Organoids can be combined with a variety of technologies to improve their functionality and range of applications. The combination of organoids and bioprinting technology allows precise control of the structure of the organoid and the creation of tissues with mechanical properties specific to the microenvironment[47]. The combination of organoids and microfluidics[48] provides precise control of the organoid culture environment, including pH, oxygen and nutrient supply, and waste removal. In addition, microfluidics can be used to create organoids with specific vascular structures, simulating the curvature and branching of blood vessels by varying the direction and velocity of fluids, a simulation that can aid in understanding the pathogenesis of disease. Choi et al.[49] demonstrated the utility of microfluidic cancer organoids using patient-derived microfluidic cancer organoids from pancreatic ductal adenocarcinoma to test several chemotherapeutic treatments, including glycogen synthase kinase inhibitors.

Organ chips simulate near-physiological tissue microenvironments by integrating the advantages of organ-like and microchip technologies[50], guiding stem cell behaviors and organ-like morphogenesis, and constructing multiorgan chips or even human body chips is the ultimate goal of this field. The liver islet-like organ chip mimics the human liver-islet axis and its glucose-stimulated response under physiological and pathological conditions[36]; notably, the coculture system promotes sensitive glucose-stimulated pancreatic islet secretion in pancreatic islet-like organs and increases glucose utilization in liver-like organs through glucose tolerance tests. Ronaldson-Bouchard et al. developed a multiorgan chip using artificial heart, bone, liver, and skin connected by recirculating vascular flow and were able to outline the pharmacokinetic and pharmacodynamic profile of adriamycin in humans in a 4-week culture. Combining microfluidic technology with organoid chips[51, 52] can provide a controlled biomimetic microenvironment by mimicking blood circulation and intercellular substance exchange in vivo. Rifes. et al. developed a microfluidic chip with a “Christmas tree” structure to create a gradient of Wnt signaling proteins on which human embryonic stem cells (hESCs) differentiate, thereby mimicking the process of early neural tube development. Deng et al.[53] used gene editing to repair disease-causing genes in UiPSCs from patients with early-onset RP, and retinal cells obtained by differentiation revealed the phenotype of healthy retinal cells. Using single-cell sequencing to analyze four early stages of retinal organoid differentiation, Lu et al. [54]. identified evolutionarily conserved patterns of gene expression during retinal progenitor cell maturation and identified differences in gene expression between the developing macula and periphery, as well as between different levels of cell populations. These results will inform future studies of human retinal development. During in vitro construction, synthetic biology[55, 56] is integrally merged with organoids to engineer cellular communities endowed with distinct functions, propelling the model’s evolution toward increased structural and functional intricacy. The strategic modulation of cellular communication, achieved through the deployment of optogenetic interfaces, distal and proximal signaling cues, and purpose-built cell adhesion mechanisms, is pivotal in orchestrating and refining the architecture of organoids.

Organoid culture technology has great potential and applications. However, some of the current limitations remain unresolved[57], such as the long-term stability and genetic variability of organoids in culture, the complexity of multicellular organisms, and the challenge of performing large-scale production, which needs to be urgently overcome. In addition, the cultural standards of the technology need to be harmonized, and ethical issues need to be continuously overcome and improved.

#### 4.3.4 Cluster 4. Molecular mechanism

As disease models, organoids can help researchers elucidate disease activation mechanisms at the molecular and cellular levels, helping to study disease intervention and treatment better t. The Wnt/beta-catenin and Notch signaling pathways play critical roles in regulating organoid stem cell fate determination, differentiation, and development. YAP signaling also contributes to stem cell renewal and plasticity, further enhancing tissue regeneration potential. Through organoid culture, researchers have revealed the complex roles of critical molecules such as p53, E-cadherin, and TGF-beta in cancer progression, invasion, and metastasis. The dynamic interactions of these signaling pathways, including kinase activation, phosphorylation events, receptor-ligand interactions, and receptor-ligand interactions, reveal a complex molecular picture of the molecules that drive tumorigenesis. They also help to study the DNA damage response, epigenetic modifications (e.g., methylation patterns), and the emergence of senescent or resistant phenotypes, informing the development of therapeutic strategies.

One study revealed that the application of single-cell DNA sequencing techniques to compare gene expression programs in organoid endocytosis and human fetal neocortical tissue was very similar[58], suggesting that human cortical development can be studied in organoid culture. Uzquiano et al.[59] have identified integrated single-cell transcriptomics, epigenetics, and spatial mapping of human cortical organogenesis and have identified genes with predictive human-specific roles in lineage establishment. In addition, Fleck et al.[60] also combined organoid and single-cell gene technologies to investigate the gene regulatory networks underlying human brain development, for example, the GLI3 transcription factor regulatory group in telencephalic development not only regulates the dorsoventral pattern with HES4/5 as a direct GLI3 target but also diversifies the ganglionic bulge. Using retinal organoids as a research model, Wahle et al.[61] performed quantitative measurements across different spatial scales and molecular modalities, integrating genomic data with spatially segmented nuclei into a multimodal atlas and highlighting pathways of RGC cell death and mosaic inheritance of retinal organoids contributing to the understanding of cell fate regulation. Álvarez-Varela et al.[62] used rectal cancer organoids to reveal the molecular mechanisms underlying the susceptibility to recurrence after chemotherapy by targeting the MEX3A+-resistant cell population to improve the treatment of advanced colorectal cancer. Twenty-five neuroendocrine tumor-like organ lines of gastroenteropancreatic tumors were established and comprehensively molecularly characterized by Kawasaki et al.[63] Complex knockdown of TP53 and RB1, as well as overexpression of key transcription factors, provided a genetic understanding of GEP-NEN and linked its genetics to its biological phenotype. Unprecedented opportunities for new therapies for gastrointestinal and other diseases through insights into disease etiology by corresponding organoid models. Organoid-based approaches pave the way for precision medicine interventions tailored to individual patient needs.

#### 4.3.5 Cluster 5. Regenerative Medicine

Regenerative medicine, as an emerging field of health sciences research, is dedicated to developing ex vivo healthy tissues to replace specific functionally or structurally compromised organs, achieving no immunosuppression, no complications, and reduced toxicity. Unlike existing regenerative medicine treatments, the tissues required for organoids can be collected through minimally invasive procedures such as biopsies, which makes it easier to ensure cell safety. The use of organoids generated from adult stem cells with a lower risk of tumor prevention or immune rejection is safer after transplantation than the use of other grafts, and the culture environment of the organoids allows for their large-scale expansion.

Various studies[64, 65] have shown that organoid-based regenerative medicine can be applied to the study of different diseases. The intestinal organoid model has a mature system and was first used in regenerative medicine. Avansino et al.[66] transplanted murine small intestinal adult stem cell-derived organoids were transplanted into the jejunum of recipient mice, and the organoids were able to grow new small intestinal mucosa in the transplanted area and differentiate into intestinal epithelial cells and cup cells. Fukuda et al.[67] found that cultured mouse epithelial organoids could reconstitute self-renewing epithelial cells in the colon through the intrinsic mechanism of the epithelium and maintain their properties in the gastrointestinal tract through the xenografting method. Liver transplantation is one of the hottest areas of research in regenerative medicine, and patient-derived liver-like organs are currently known to cause immune rejection after transplantation. With large-scale in vitro expansion, the use of liver-like organs is expected to solve the problems of limited graft sources and a shortage of quantity. HU et al.[68] transplanted human fetal liver-like organs into a mouse model of liver injury, and the results also demonstrated that liver-like organs can significantly expand and regenerate damaged livers after transplantation.

Organoids are combined with gene editing and nanotechnologies to provide new ideas for the treatment of diseases such as chronic kidney disease, biliary atresia, inflammatory bowel disease, and metabolic liver disease. Hans Clevers et al.[69] used CRISPR-dependent single-base editing to edit and repair mutations in the CFTR causative gene in cystic fibrosis patient samples from the Dutch Organoid Biobank, providing a clinically matched therapeutic option for cystic fibrosis disease. Wu et al. established a multifunctional blastocyst complementation platform based on CRISPR-Cas9-mediated genome editing of fertilized eggs. The rat pancreas was grown in vivo in mice, and interspecies blastocyst complementation was successfully demonstrated in pancreas knockout mice. Personalized stem cell organoids can overcome the immune rejection of artificial organ transplants, and the deep integration of micro- and nanoengineering-based organ maturation and personalized organoid transplantation can lead to regenerative organ therapy. Nanotechnology-constructed biomaterial scaffolds provide an environment conducive to cell adhesion, differentiation, and proliferation for the self-organization of organoid pluripotent stem cells[70]. In addition, microfluidic technology controls the differentiation of stem cells to repair nervous system damage, providing new ideas for regenerative medicine.

Big data, cross-fertilization of disciplines, and the wide application of technologies such as life histology, gene editing, high-resolution imaging, and new material manufacturing in the field of regenerative medicine have pushed the breadth and depth of regenerative medicine research to continue expanding, and the scope of the field is constantly broadening, which has also brought new opportunities for the development of regenerative medicine.

## Limitations

As with other bibliometric studies, this paper has several limitations. Only English-language publications were selected, and there may be language bias. There are differences in authors’ and institutions’ attributions, which may cause bias in the statistical results.

## Conclusion

Organoids, a breakthrough in the field of stem cell research in recent years, provide medicine with a powerful tool for the study of human development and disease modeling. As organoid research continues, organoids will play an increasingly important role in the medical field and scientific research. Influential authors and institutions, collaborative networks, and emerging research themes also provide invaluable resources for subsequent researchers and policymakers to further advance the field’s exploration.

